# Stable, multigenerational transmission of the bean seed microbiome despite abiotic stress

**DOI:** 10.1101/2024.03.21.586100

**Authors:** Abby Sulesky-Grieb, Marie Simonin, A. Fina Bintarti, Brice Marolleau, Matthieu Barret, Ashley Shade

## Abstract

Seed microbiomes initiate plant microbiome assembly, but the consequences of environmental conditions of the parent plant for seed microbiome assembly and transmission are unknown. We tracked endophytic seed bacterial communities of common bean lines exposed to drought or excess nutrients, and discovered stable transmission of 22 bacterial members regardless of parental plant treatment. This study provides insights into the maintenance of plant microbiomes across generations, even under challenging environmental stress.

## Main Text

The seed microbiome plays a role in both pathogen transmission and in shaping a plant’s beneficial microbiome^1,2^. Understanding the inheritance of microbiome members over plant generations, especially as mediated through seed transmission, is of interest for plant microbiome management and conservation^3–7^. Plant microbiomes can be assembled via both horizontal transmission from the environment and vertical transmission from the parent plant through the seed microbiome^8–10^. For seed transmission, microbial cells can either migrate through the vascular tissue or floral compartments and become packaged within the internal seed tissues or colonize seed surfaces^10–14^. During germination, seed microbiome members that first colonize the plant can determine priority effects and drive the trajectory of microbiome assembly^15–17^. Additionally, these “inherited” microbiome members enriched by the parent plant may provide important benefits for the offspring ^18–21^.

A remaining unknown is how the environmental conditions of the parent plant influence seed microbiome assembly and whether stress can drive long-term alterations in the microbiome across generations^22–24^. A few studies have found an influence of parental plant line on the resulting seed microbiome of the next generation^5,25^. Other studies have suggested an impact of parental stress on the seed microbiome after one generation, particularly drought in beans and wheat^26,27^, excess nutrients in beans^26^, and salt stress in rice^28^. A legacy effect of environmental stress on the seed microbiome could have consequences for the healthy assembly of the microbiome in the next plant generation^29^.

We conducted a multigenerational experiment in which we exposed common bean plants (*Phaseolus vulgaris L.*) to drought or high nutrients during their early vegetative growth (**Fig. S1**). Plants were grown in agricultural soil in environmental chambers to allow for the natural assembly of their native microbiome while also controlling environmental conditions. We were motivated to study *Phaseolus vulgaris* L. because it is a critical legume for global food security, supporting the health and livelihood of millions of people worldwide^30,31^. We chose these two abiotic treatments, drought, and nutrient excess, because of their relevance to bean production in a changing climate. Bean production is threatened by drought associated with warming and changes in precipitation patterns^32–34^. Furthermore, beans grown in areas of high production are often managed with excess mineral fertilizer, which can impact the selection and stability of the plant microbiome^35^.

We hypothesized that abiotic treatment of the parent plant characteristically alters the seed endophyte microbiome. Furthermore, we expected that each abiotic treatment would have specific consequences for the composition and stability of taxa transmitted across generations. We tested our hypothesis by planting a starting set of G0 bean seeds through two generations, exposed to either control growth conditions, drought, or excess nutrient treatments while tracking each plant’s parental line within a fully factorial design (**Fig. S1**). We applied standard 16S V4 rRNA gene amplicon sequencing of the seed bacterial endophyte communities for each of the three generations to assess the impact of the treatments on the seed microbiome and the potential for transmission of the bacterial taxa.

The drought and nutrient treatments did not influence microbiome alpha or beta diversity in either generation (**Fig. S2, Fig. S3, Table S1**). Furthermore, we detected no legacy influence of the G1 treatments on the G2 seed microbiomes and no overarching differences in the microbiomes across the G1 and G2 generations (**Fig. S3**, **Table S1**). However, there was an appreciable influence of parental line for those G2 plants that originated from the G1 drought treatment (**Fig. S4B**, PERMANOVA *post-hoc* r^2^=0.40311 F=1.4637 p=0.0021), but not for the control or nutrient parental lines.

Instead, we found evidence of stable transmission of 128 of 658 detected Amplicon Sequence Variants (ASVs) across all three generations, regardless of the experimental treatment (**Fig. 1A**). The 128 ASVs detected in all three generations were in higher relative abundance than the ASVs that were only seen in one or two generations (Kruskal-Wallis test: test-statistic=363.59, df=2, p<0.0001; Post-hoc Dunn’s test with Benjamini-Hochberg correction: 1 v. 3 generations: test-statistic=17.78, adjusted p<0.0001, 2 v. 3 generations: test-statistic=5.879, adjusted p<0.0001) (**Fig. 1B**). Furthermore, the Genus-level taxonomic profiles of G0, G1, and G2 microbiomes were highly comparable (**Fig. S5)**, suggesting a consistent taxonomic signature of the ASVs detected across seed generations.

**Figure 1.**
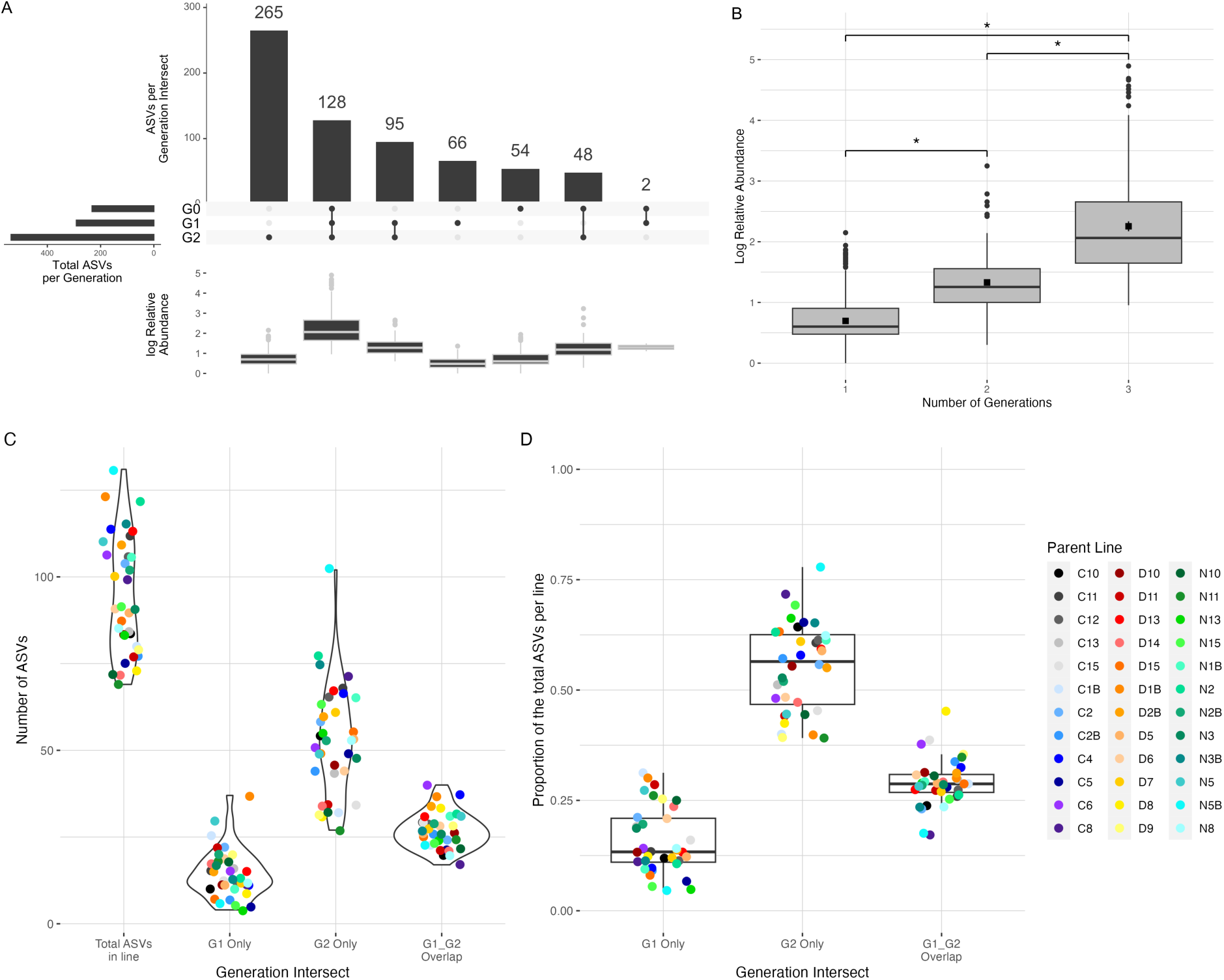
Unique and overlapping ASVs between generations. **A,** Number of ASVs and relative abundance of ASVs per generation intersect across all samples. G0 n=20 samples (five seeds per sample), G1 n =36, G2 n=108. **B,** Log relative abundance of ASVs based on how many generations in which they are found. Out of 658 total ASVs detected, ASVs found in all three generations are significantly more abundant in the dataset than ASVs found in only one or two generations. Black squares indicate the mean value. (Kruskal-Wallis test: test-statistic=363.59, df=2, p-value<0.0001; Post-hoc Dunn’s test with Benjamini-Hochberg correction, 1 vs 3 generations: test-statistic=17.78, adjusted p<0.0001, 2 vs 3 generations: test-statistic=5.879, adjusted p<0.0001). **C,** Total number of ASVs per parent line and number of ASVs found in G1, G2, or overlapping. “G1_G2 Overlap” is defined as ASVs present in both the G1 sample and at least one G2 offspring within a parent line. 99 ASVs were identified as overlapping within parent lines, and there were overlapping ASVs identified in all 36 lines. **D,** proportion of the total ASVs per line found in G1, G2, or overlapping. Boxplots represent the median values and first and third quartiles, and whiskers represent the 95% confidence interval. C, D, and N in parent line IDs denote lines that received Control, Drought or Nutrient treatment in G1, respectively.

We identified ASVs overlapping between a G1 parent plant and at least one of its offspring in G2, which we call “overlapping ASVs.” There were 99 overlapping ASVs discovered among the 36 lines, 70 of which were found in at least two parent lines and 43 of which were common to all treatments. The proportion of overlapping ASVs in each line ranged from 17% to 45% of the total ASVs detected (**Fig. 1C, 1D**).

Nine overlapping ASVs were present in all 36 parental lines, and an additional 13 ASVs were found in at least half of all parental lines. These 22 ASVs generally had 100% transmission to all three G2 offspring per parent line (**Fig. 2**). At the same time, we noticed that less prevalent ASVs were not consistently transmitted in all offspring within lines. The G1 treatment did not impact the average transmission of the ASVs in the G2 offspring, and there was no significant difference in average transmission between parental lines (Pearson’s Chi-squared Test. G1 Treatment: X^2^=0.67413, df=4, p=0.9545. Line: X^2^=63.48, df=70, p=0.6958) (**Fig. 2**).

**Figure 2.**
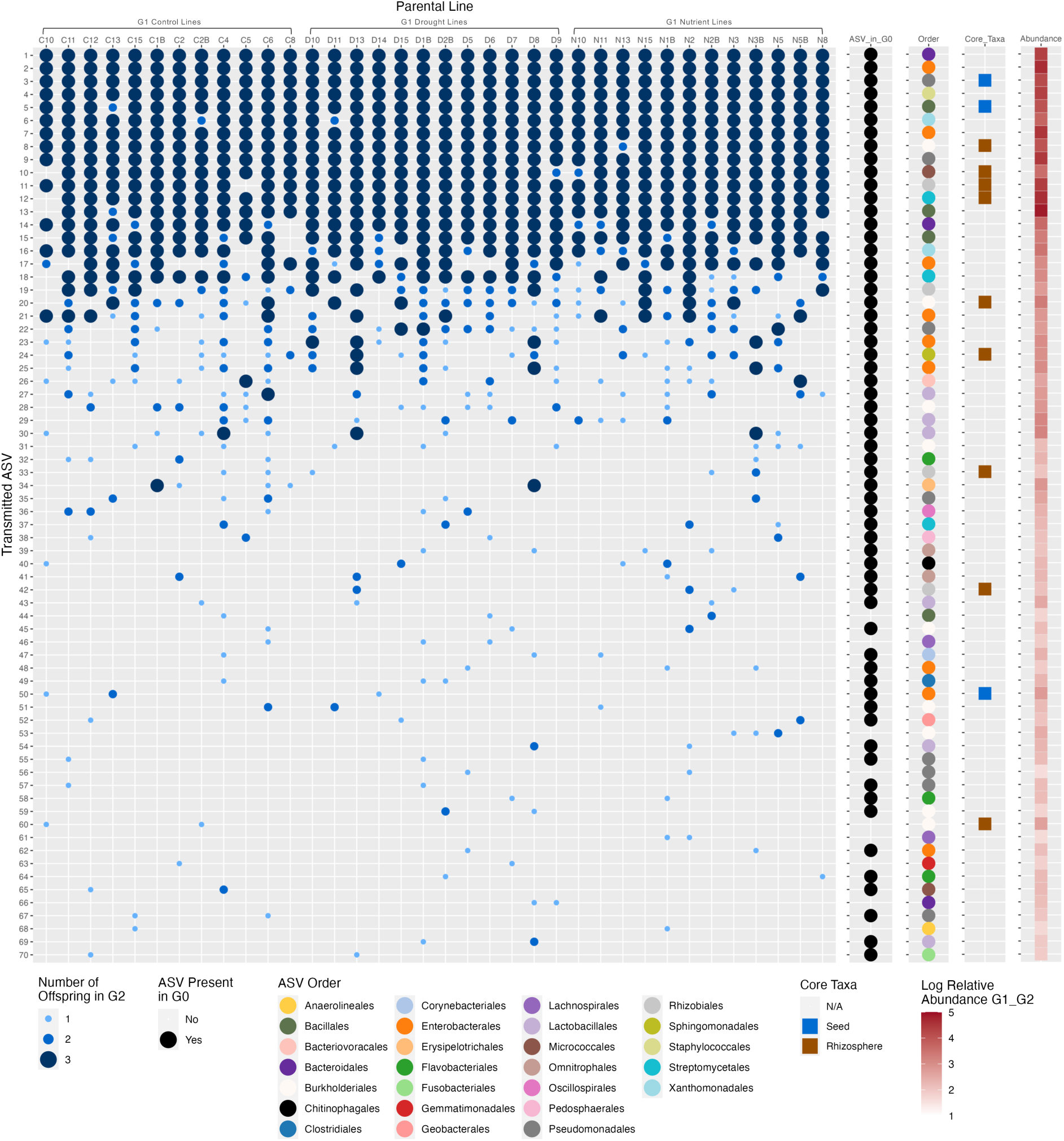
Prevalence in lines, transmission, core taxa identity and relative abundance of overlapping ASVs in G2 offspring. Of the 99 ASVs that were found overlapping between G1 and G2 within parent lines, 29 ASVs that were only in one parent line were removed, and the remaining 70 ASVs are listed above, ordered from presence in the highest number of lines to lowest number of lines. Blue dots represent the number of G2 offspring containing the ASV in each line. 61 of these ASVs are also found in the G0 dataset indicated by black dots. The taxonomy of each ASV identified at the Order level is indicated by colored dots in the Order column (26 Orders). Blue and brown squares in the Core_Taxa column indicate identity with seed or rhizosphere core taxa, respectively. Red boxes in the right-most column represent the log relative abundance of the ASV overall across G1 and G2. There is no significant difference in ASV transmission in G2 offspring between G1 treatments or parental lines (Pearson’s Chi-squared Test. G1 Treatment: X^2^=0.67413, df=4, p=0.9545. Line: X^2^=63.48, df=70, p=0.6958).

Most overlapping ASVs that were found in at least two parental lines were also detected in the G0 microbiomes (61 out of 70 ASVs) (**Fig. 2**). These ASVs were taxonomically diverse and included 39 Families and 26 Orders. Many overlapping ASVs were also highly abundant (**Fig. 2**). Their Genus-level taxonomic compositions in G2 were very similar within and across parent lines (**Fig. S7**).

To seek generalities with other relevant studies, we compared the 70 overlapping ASVs detected in this study to the six “core” common bean seed microbiome ASVs identified by Simonin *et al.* 2022^36^ and to the 48 “core” common bean rhizosphere OTUs identified by Stopnisek and Shade 2021^37^. These previously reported core taxa were identified across multiple studies and are hypothesized to be important for health in common beans. Three core common bean seed ASVs^36^ were found in the overlapping G1-G2 dataset, specifically from the *Pseudomonas*, *Bacillus*, and *Pantoea* genera (**Table 1**, Fig. 2, **Fig. S6**). Two ASVs, the *Pseudomonas* and *Bacillus* core members, were detected in all 36 parental lines, while the *Pantoea* core seed microbiome member was found in only three parental lines (**Table 1**). There were nine ASVs aligned at >96% identity to the previously identified core rhizosphere taxa^37^ (**Table 1, Table S3, Fig. 2**). These ASVs were classified in the families Comamonadaceae, Devosiaceae (genus *Devosia*), Methyloligellaceae, Oxalobacteraceae (genus *Massilia*), Rhizobiaceae (genus *Ochrobactrum*), Sphingomonadaceae (genus *Sphingomonas*), Streptomycetaceae (genus *Streptomyces*), and Micrococcaceae in the genus *Arthrobacter*, of which our seed ASV aligned at 98.8% identity to the most abundant core OTU in the rhizosphere study (**Table 1, Table S3**). These genera are commonly associated with plant microbiomes^38–42^.

**Table 1.**
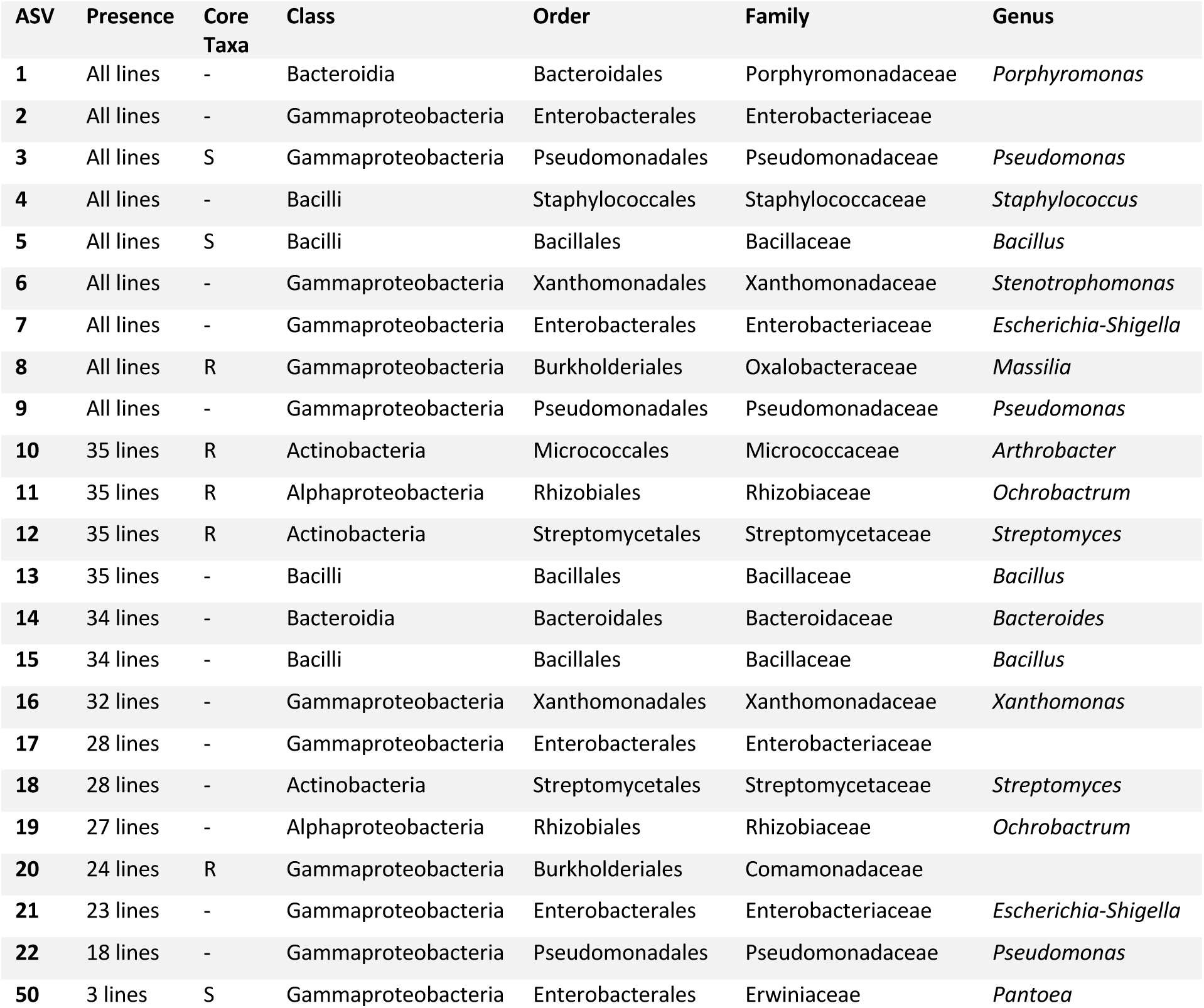
Stable ASVs that were detected in all three generations. Nine ASVs are found in all 36 parental lines, two of which are core seed microbiome taxa, labeled “S”. 13 additional ASVs were present in 50% or more of the parental lines. An additional core seed microbiome member was found in three parental lines. Five of these ASVs align to bean rhizosphere core OTUs at >96% identity, labeled “R”. All ASVs listed are also found in Generation 0 seeds.

| ASV | Presence | Core Taxa | Class | Order | Family | Genus |
| --- | --- | --- | --- | --- | --- | --- |
| 1 | All lines | - | Bacteroidia | Bacteroidales | Porphyromonadaceae | *Porphyromonas* |
| 2 | All lines | - | Gammaproteobacteria | Enterobacterales | Enterobacteriaceae |  |
| 3 | All lines | S | Gammaproteobacteria | Pseudomonadales | Pseudomonadaceae | *Pseudomonas* |
| 4 | All lines | - | Bacilli | Staphylococcales | Staphylococcaceae | *Staphylococcus* |
| 5 | All lines | S | Bacilli | Bacillales | Bacillaceae | *Bacillus* |
| 6 | All lines | - | Gammaproteobacteria | Xanthomonadales | Xanthomonadaceae | *Stenotrophomonas* |
| 7 | All lines | - | Gammaproteobacteria | Enterobacterales | Enterobacteriaceae | *Escherichia-Shigella* |
| 8 | All lines | R | Gammaproteobacteria | Burkholderiales | Oxalobacteraceae | *Massilia* |
| 9 | All lines | - | Gammaproteobacteria | Pseudomonadales | Pseudomonadaceae | *Pseudomonas* |
| 10 | 35 lines | R | Actinobacteria | Micrococcales | Micrococcaceae | *Arthrobacter* |
| 11 | 35 lines | R | Alphaproteobacteria | Rhizobiales | Rhizobiaceae | *Ochrobactrum* |
| 12 | 35 lines | R | Actinobacteria | Streptomycetales | Streptomycetaceae | *Streptomyces* |
| 13 | 35 lines | - | Bacilli | Bacillales | Bacillaceae | *Bacillus* |
| 14 | 34 lines | - | Bacteroidia | Bacteroidales | Bacteroidaceae | *Bacteroides* |
| 15 | 34 lines | - | Bacilli | Bacillales | Bacillaceae | *Bacillus* |
| 16 | 32 lines | - | Gammaproteobacteria | Xanthomonadales | Xanthomonadaceae | *Xanthomonas* |
| 17 | 28 lines | - | Gammaproteobacteria | Enterobacterales | Enterobacteriaceae |  |
| 18 | 28 lines | - | Actinobacteria | Streptomycetales | Streptomycetaceae | *Streptomyces* |
| 19 | 27 lines | - | Alphaproteobacteria | Rhizobiales | Rhizobiaceae | *Ochrobactrum* |
| 20 | 24 lines | R | Gammaproteobacteria | Burkholderiales | Comamonadaceae |  |
| 21 | 23 lines | - | Gammaproteobacteria | Enterobacterales | Enterobacteriaceae | *Escherichia-Shigella* |
| 22 | 18 lines | - | Gammaproteobacteria | Pseudomonadales | Pseudomonadaceae | *Pseudomonas* |
| 50 | 3 lines | S | Gammaproteobacteria | Enterobacterales | Erwiniaceae | *Pantoea* |

Our multigenerational study indicates that numerous seed microbiome members were consistently packaged in the seed endophytes of the G0, G1, and G2 plants, regardless of the growth conditions (field or environmental chamber), the starting soil or abiotic treatment applied to the plants. This suggests a consistent and stable seed microbiome transmission for common beans. This is an unexpected result because, for many plant species, the seed microbiome has been reported to have relatively low diversity (tens to hundreds of taxa), low biomass/small community size (dozens of cells), and notably high compositional variability^4,36,43^, suggesting an influence of stochasticity in the assembly. Our results indicate that against a background of high variability, a handful of members of the seed endophyte microbiome exhibit stable transmission. These transmitted taxa could not have been identified before, because previous studies have focused on only one generation, and, perhaps more importantly, on seeds that were polled across multiple parent plants. These stable seed microbiome taxa may establish mutualistic or commensal associations and persist through the plant life cycle until packaged within the seed for the next generation. It also suggests that a parental effect on the seed microbiome has the potential to outweigh abiotic effects.

Our results cannot address the exact mode of transmission of the seed endophyte microbiome, which is a limitation of the work. It remains unclear whether the seed endophytes are passed directly *via* vertical transmission (e.g., from seed to plant to seed via the vascular tissue as previously reported as the primary pathway for common bean^11,44^) or re-acquisition (e.g., parent plants selectively recruit the same taxa from the environment). Regardless of the transmission mode, the seed microbiome members’ stability despite abiotic treatment, their prevalence and high relative abundance, and their consistent detection across several bean microbiome investigations suggest a strong selection of the plant seed environment for these taxa. Thus, while our initial hypothesis about the importance of abiotic treatment in driving seed endophyte microbiome variation was not supported, there was clear evidence that a stable seed microbiome was transmitted in all conditions in our study.

As the need for sustainable solutions to maintain or improve agricultural productivity increases, plant microbiome management, microbiome engineering, and breeding plants for improved microbiomes will be critical strategies^45–47^. Applying beneficial plant microbiome members via seed treatments or soil inoculation has shown promise in improving plant growth or health^48,49^. The stably transmitted bean seed microbiome members identified here provide targets for future research to understand how to shape legume microbiomes to improve crop yield, health, and resilience^50,51^.

## Acknowledgments

This work was supported by the United States Department of Agriculture award 2019-67019-29305 to AS and MB and by the Michigan State University Plant Resilience Institute. AS acknowledges support from the United States Department of Agriculture National Institute of Food and Agriculture and Michigan State University AgBioResearch, and the Centre National de la Recherche Scientifique (CNRS), France.

## Online Methods

### Bean cultivar

*Phaseolus vulgaris L.* var. Red Hawk^52^, developed by the Michigan State University Bean Breeding program, was selected as a representative dry bean crop. Red Hawk seeds were obtained from the Michigan State University Bean Breeding Program from their 2019 harvest and stored at 4°C until ready for use in experiments. These seeds were obtained as “Generation 0” and used to plant the first generation of the experiment.

### Soil preparation

Agricultural field soil was collected for each planting group in September 2019, December 2020, May 2021, and September 2021 from a Michigan State University Agronomy Farm field that was growing common beans in 2019 (42°42’57.4”N, 84°27’58.9”W, East Lansing, MI, USA). The soil was a sandy loam with an average pH of 7.2. When collecting the soil, we avoided the dry top layer of soil and plant debris. Field soil was covered with air-tight lids and stored at 4°C until the experiment began. The soil was passed through a 4mm sieve to remove rocks and plant debris and then mixed with autoclaved coarse vermiculite at a 50% v/v ratio.

### Surface sterilization and seed germination

Red Hawk seeds were surface sterilized prior with a solution of 10% bleach and 0.1% Tween20. Seeds were randomly selected from the bulk G0 seed supply or the harvested Generation 1 (G1) seed supply, but we avoided seeds that were visibly cracked or moldy.

Approximately 20 seeds were placed in a sterile 50 mL conical tube, 20-30 mL of bleach solution was added, and then the seeds were soaked for 10 minutes, with agitation at 5 minutes. After soaking, the seeds were rinsed 5 times with sterile DI H_2_O. On the final rinse, 100 μL of rinse water was spread onto Tryptic Soy Agar (TSA) and Potato Dextrose Agar (PDA) plates to assess the efficacy of the seed surface sterilization. TSA plates were incubated overnight at 28°C and PDA plates at room temperature for 48 hours. Seeds corresponding to plates that had microbial growth were discarded from the experiment and replaced with surface-sterile ones. For germination, seeds were placed in a Petri dish lined with sterile filter paper and supplemented with 1-2 mL sterile DI H_2_O. Petri dishes were stored in the dark at room temperature for 3-4 days, with an additional 2 mL sterile DI H_2_O added after two days. Once seeds had sprouted radicle roots, they were transferred to soil.

### Growth conditions

For G1, three germinated seeds were planted into 3.78-liter pots filled with the soil-vermiculite mixture and placed in a high-light BioChambers FLEX™ LED growth chamber with a 16-hour day/8-hour night cycle at 26°C and 22°C, respectively, and 50% relative humidity. Once seedlings emerged and reached the VC growth stage (vegetative growth with two cotyledons and primary leaves expanded), they were thinned to one seedling per pot. Plants were watered every other day with 300 mL 0.05% 15-10-30 water-soluble fertilizer solution (control condition) (Masterblend International, Morris, IL, USA). At the V3 stage (vegetative growth with third trifoliate leaves expanded), treatments began for the drought- and nutrient-treated plants. Drought plants received 100 mL of 0.15% 15-10-30 fertilizer solution every other day (66% less water than control with the same concentration of nutrients), and nutrient plants received 300 mL of 0.15% 15-10-30 fertilizer solution every other day (3X concentrated nutrients with the same volume of water as control). After approximately 14 days of the treatment period, when plants reached the R1 stage (reproductive stage, first open flowers), they were returned to the water regime for the control plants (every other day) until senescence. As plants began to dry, they were watered less frequently as needed. Mature seeds were collected from 12 plants per treatment for seed microbiome assessment and for germination for the next generation.

For Generation 2 (G2), seeds from the 36 G1 parental lines that received either control, drought, or nutrient conditions were planted in a full factorial design and grown under one of the three treatment conditions in G2. There were nine cross-generational treatment combinations total (G1_G2, n=12 plants per treatment): Control_Control, Control_Drought, and Control_Nutrient; Drought_Control, Drought_Drought, and Drought_Nutrient; Nutrient_Control, Nutrient_Drought, and Nutrient_Nutrient. Six seeds from each parental line were surface sterilized and germinated as described above, then planted in the field soil-vermiculite mixture in seedling trays in the growth chamber under the conditions stated above. G2 plants were grown in three randomized planting groups, with each planting group containing parental lines from all three treatments. Once plants reached the VC stage, three healthy seedlings per parent line were transferred to 3.78-liter pots. Each G1 parental line provided one offspring per G2 treatment for a total of 108 plants in G2. Plants were watered according to the conditions and treatment timeline in G1.

### Seed harvest

Once the plants had senesced and pods were dried, seeds were harvested for planting or microbiome analysis. Seed pods were removed from each plant and stored in sterile Whirlpak bags. Pods and seeds per plant were counted, and then seeds were removed from the pods and pooled by the parent plant in 50 mL conical tubes and stored at 4°C for use in planting. Five seeds per plant were selected for microbiome analysis and stored in a 15 mL conical tube at −80°C until DNA extraction was performed.

### DNA extractions

DNA extractions were performed on sets of five randomly selected seeds per plant, which is our unit of microbiome sampling. For the G0 bulk seed, twenty sets of five randomly selected seeds from each plant were analyzed (these five seeds did not necessarily come from the same parent plant). Seeds were analyzed from 36 parent plants for G1 and from 108 offspring for G2. Seeds were thawed and surface sterilized according to the method above, and then microbial DNA was extracted from the endophytic compartment using a protocol adapted from Barret *et al.* 2015 and Bintarti *et al.* 2021^8,43^. Following surface sterilization, the seeds were sliced in half lengthwise along the natural division of the cotyledons with a sterile razor blade. Sliced seeds were placed in a 50 mL conical tube, and 20-30 mL of sterile Phosphate-buffered Saline (PBS) with 0.05% Tween 20 was added. Seeds were soaked overnight at 4°C with constant agitation on a surface shaker at 160 rpm. After soaking, tubes were centrifuged at 4500xg, 4°C, for one hour. Seed tissue and supernatant were removed, and the remaining pellet was transferred to a 1.5 mL microcentrifuge tube. Pellets were stored at −80°C until extraction with the E.Z.N.A. Bacterial DNA kit (Omega Bio-tek, Inc., Norcross, GA, USA) following the manufacturer’s protocol with the following modifications. The seed material pellet was resuspended in 100 μL TE Buffer, 10 μL kit-provided Lysozyme was added, and the samples were vortexed thoroughly and incubated at 37°C for 1 hour. The glass bead step from the

E.Z.N.A. kit was utilized with 25-30 mg glass beads provided, and samples were vortexed at maximum speed for 10 minutes in a 24-tube vortex adapter. After adding the Proteinase K, the samples were incubated in a shaking heat block at 55°C for 2 hours. In the final step, DNA was eluted in 60 μL Elution Buffer and incubated at 65°C for 10 minutes before centrifuging into the final tube.

DNA extractions were performed in randomized batches within each generation (**Table S2**). For each batch, negative and positive controls were included. The negative control was 3 mL sterile PBS+Tween buffer, and the positive control was an aliquot of a mixture of cells from a custom-made mock bacterial community in 3 mL buffer^53^. These controls were soaked overnight alongside the seed samples and then processed and sequenced as described for the seeds, and then ultimately used to perform batch-informed bioinformatic sequence decontamination^54^.

### Amplicon Sequencing

Sequencing of the V4 region of the 16S rRNA gene (515F-806R)^55,56^ was performed at the Environmental Sample Preparation and Sequencing Facility (ESPSF) at Argonne National Laboratory (Lemont, IL, USA). The DNA was PCR amplified with region-specific primers that include sequencer adapter sequences used in the Illumina Nextseq2K flowcell; FWD:GTGYCAGCMGCCGCGGTAA; REV:GGACTACNVGGGTWTCTAAT^55–59^. Each 25 µL PCR reaction contained 9.5 µL of MO BIO PCR Water (Certified DNA-Free), 12.5 µL of QuantaBio’s AccuStart II PCR ToughMix (2x concentration, 1x final), 1 µL Golay barcode tagged Forward Primer (5 µM concentration, 200 pM final), 1 µL Reverse Primer (5 µM concentration, 200 pM final), and 1 µL of template DNA. The conditions for PCR were as follows: 94 °C for 3 minutes to denature the DNA, with 35 cycles at 94 °C for 45 s, 50 °C for 60 s, and 72 °C for 90 s, with a final extension of 10 min at 72 °C to ensure complete amplification. Amplicons were then quantified using PicoGreen (Invitrogen) and a plate reader (Infinite® 200 PRO, Tecan). Once quantified, the volumes of each of the products were pooled into a single tube in equimolar amounts. This pool was then cleaned up using AMPure XP Beads (Beckman Coulter) and then quantified using a fluorometer (Qubit, Invitrogen). After quantification, the pool was diluted to 2 nM, denatured, and then diluted to a final concentration of 6.75 pM with a 10% PhiX spike for sequencing.

Amplicons were sequenced on a 251bp x 12bp x 251bp NextSeq2000.

### Sequence data processing

Fastq files were processed in QIIME2 after primer removal by the sequencing center (QIIME2 version: 2022.8.0)^60^. Sample fastq files were imported to QIIME2 format, and samples were denoised, truncated, and merged using DADA2 with a forward truncation length of 191 and reverse truncation length of 84^61^. Amplicon sequence variants (ASVs) were defined at 100% sequence identity, and 16S taxonomy was assigned with the Silva database release 138 at the default confidence value of 0.7, and taxonomy and ASV tables were exported for further analysis in R^62^.

Data analyses were performed in R version 4.3.1 and R Studio version 2023.06.1+524^63^. There were 126.8 million merged DNA reads prior to host removal and decontamination. ASV, taxonomy, metadata tables, and phylogenetic tree files were imported into the phyloseq package, and host reads classified as chloroplast and mitochondria were removed using the subset_taxa() command in the phyloseq package version 1.44.0^64^. 90% of the total DNA reads, and 13% of the ASVs were removed as host reads, leaving 12.3 million total bacterial DNA reads. Datasets were decontaminated with the decontam package version 1.20.0 at the 0.1 threshold using the negative and positive controls from each extraction group^65^. After decontamination, there were 422,719 total DNA reads with a range of 456-5788 reads per sample in the full dataset (**Fig. S8**). Rarefaction curves were created using the rarecurve() command in the vegan package version 2.6-4^66^. Datasets were then subset for further analysis using the ps_filter() command in the microViz package version 0.10.10^67^. Since seed microbiomes typically have low bacterial diversity containing tens to hundreds of taxa, and vertical transmission of specific ASVs was a primary area of investigation in this study, the full dataset was preserved to ensure full observation ASVs^43^.

### Ecological Analysis

Alpha diversity was assessed in R using estimate_richness() in phyloseq with an ANOVA, and figures were created using the plot_richness() command from phyloseq with the ggplot2 package version 3.4.2^68^. Faith’s Phylogenetic Diversity was calculated with calculatePD() from the biomeUtils package version 0.022^69^. ANOVAs were performed with the base R stats command aov(). Weighted UniFrac distances were calculated with distance() in phyloseq and used for all analyses of beta diversity, and PERMANOVA statistical tests were performed with adonis2() from the vegan package. We used Weighted UniFrac distance because it explained the most microbiome variation relative to other resemblances we also considered (e.g., Bray-Curtis, Jaccard, etc). Post-hoc analysis on the PERMANOVA results was performed with pairwise.adonis2() from pairwise Adonis version 0.4.1^70^. Beta dispersion was assessed with the betadisper() and permutest() commands from the vegan package. **Figure 1A** was created with the UpSetR package version 1.4.0^71^ and statistical analyses were performed with leveneTest() from the car package^72^ and kruskal_test() and dunn_test() from the rstatix package^73^. Beta diversity ordinations were created with ordinate() from phyloseq with ggplot2. Additional data analysis was performed in the tidyverse package version 2.0.0^74^ and dplyr package version 1.1.2^75^. Amplicon sequence variant (ASV) transmission was analyzed as count data and Pearson’s Chi-squared Tests were performed with chisq.test() from the base R stats package. Seed core microbiota identified in Simonin *et al.* 2022^36^ were compared to transmitted ASVs, and Venn Diagrams were produced with the VennDiagram package version 1.7.3^76^. To compare the prevalent ASVs in this study to the 48 core bean rhizosphere microbiome taxa identified by Stopnisek and Shade 2021^37^, the fasta sequences for each core OTU were used as a query set in a two sequence nucleotide BLAST on the National Center for Biotechnology Information (NCBI) database website, and the fasta sequences from the seed ASVs were compared to the 48 core taxa at >96% identity^77^.

### Data Availability

Data analysis code can be found at (https://github.com/ShadeLab/Seed_transmission_Common_Bean). Raw sequences can be found on the NCBI Sequence Read Archive under BioProject number PRJNA1058980.

## Supplemental Data

**Fig. S1.**
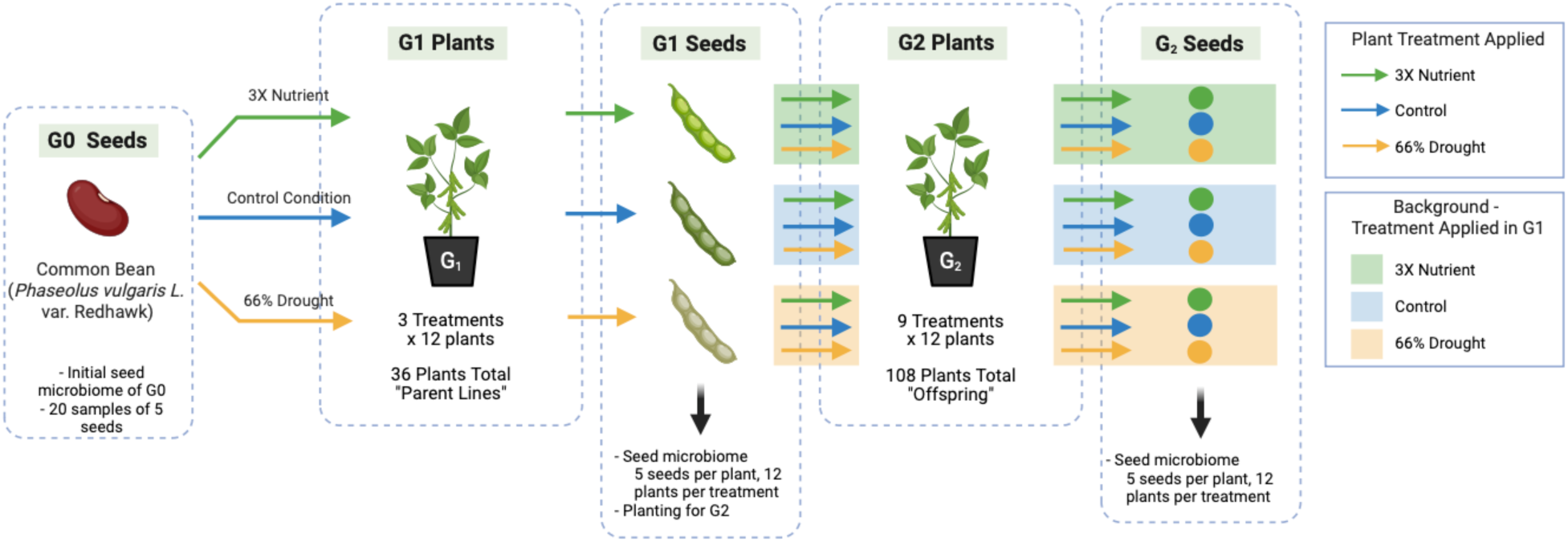
Experimental Design. Seed microbiome samples were taken from the G0 seed pool and G1 and G2 plants. Treatments were applied in G2 in a full factorial design, where one offspring from each G1 parent line was treated with each of the three treatments in G2.

**Fig. S2.**
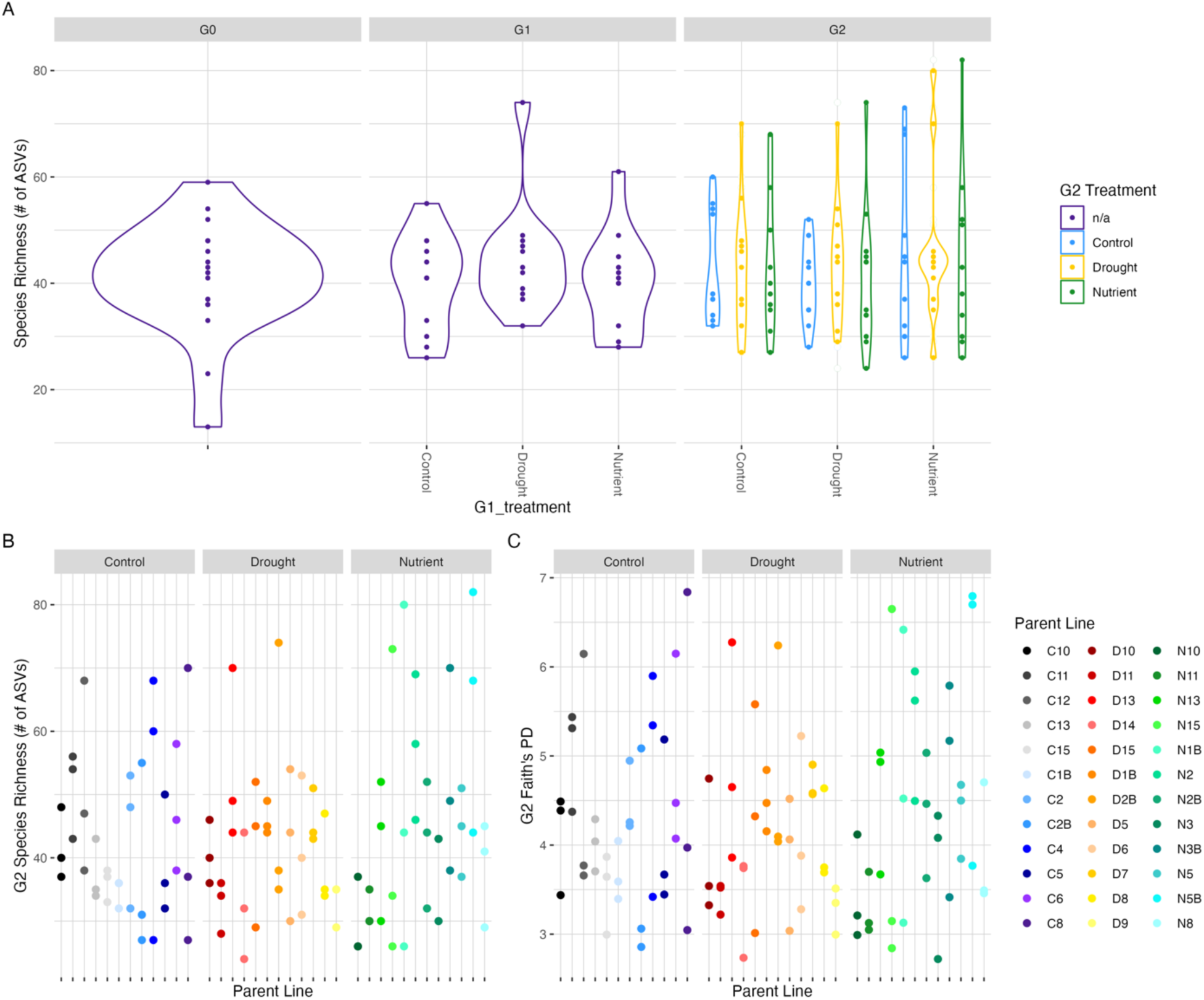
Alpha Diversity. **(A)** Number of ASVs observed in each of the seed microbiome samples across all generations. Five seeds were used for each sample. G0 n=20 samples, G1 and G2 n=12 samples per treatment group. There is no influence of treatment groups on the species richness observed in either G1 or G2 (ANOVA, G1_treatment: F= 0.150, p-value=0.861. G1_G2: F=0.393, p-value=0.923). **(B)** Number of ASVs observed, and **(C)** Faith’s Phylogenetic Diversity of G2 seed samples by parental line. Gray bars indicate treatment applied to the parent plant in G1. There are no significant differences between parent lines in either Richness or PD measure (ANOVA, Richness: F=1.122, p=0.334; PD: F=1.111, p=0.346).

**Fig. S3.**
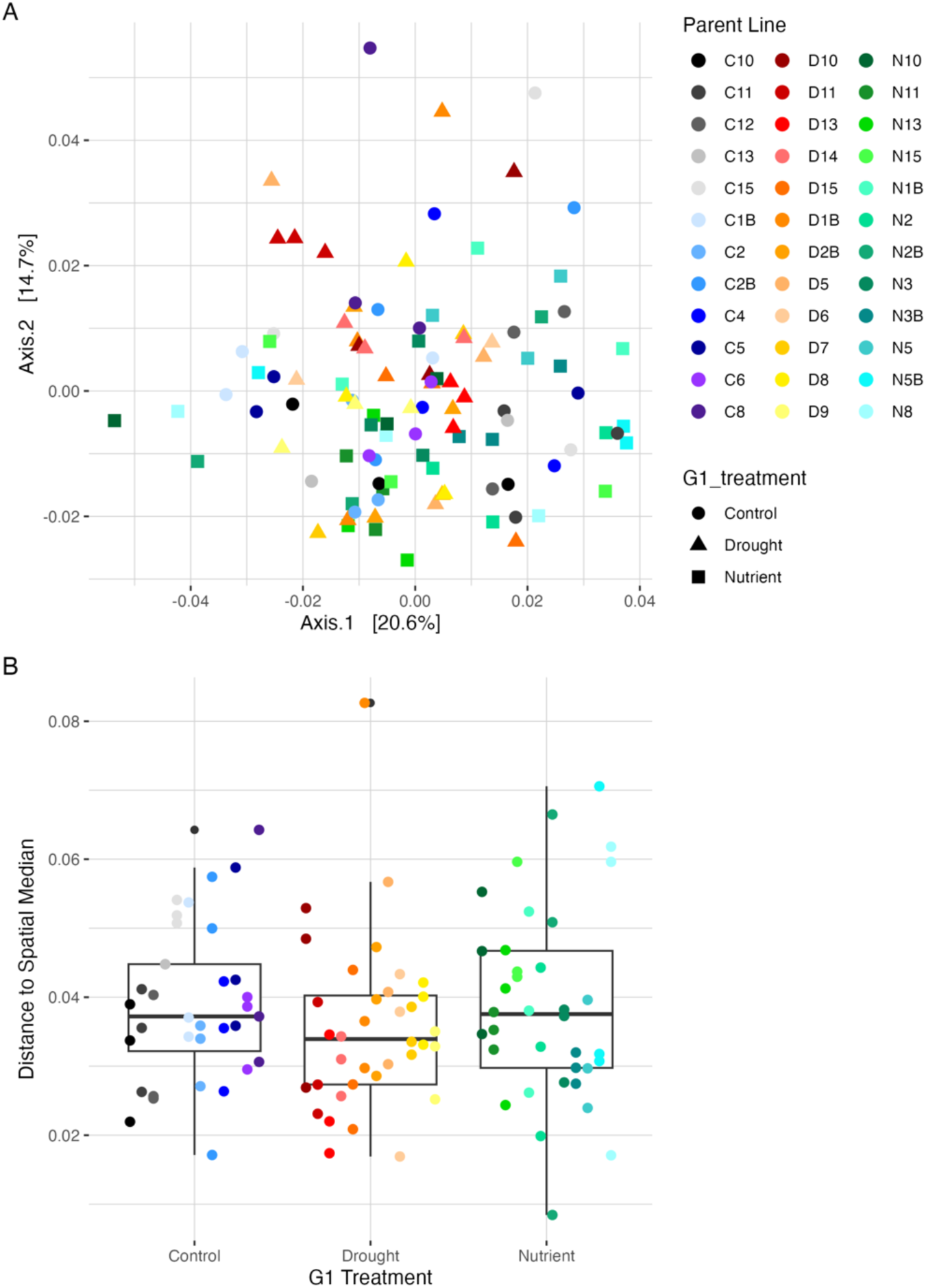
Beta diversity in Generation 2 seed samples. **(A)** PCoA of Weighted Unifrac distance of G2 seed samples. Points represent three offspring from each parent line, each of which received a different treatment in G2. Parent plant line is the only significant explanatory variable in the G2 samples (PERMANOVA, r^2^=0.356, F= 1.2421, p=0.0065**). **(B)** Beta dispersion around the spatial median of Weighted Unifrac distances in G2 seed samples. Lines are grouped by G1 parent treatment, represented by black boxplots. G1 treatment and parent line are not significant. (ANOVA, G1 Treatment: DF=2, F-value= 1.2835, p= 0.2835. Line: DF: 35, F-value=1.0825, p=0.3714). Sample G2_9, the line C13 Nutrient offspring, was removed from the figures as an outlier. However, statistics were performed with this sample included.

**Fig. S4.**
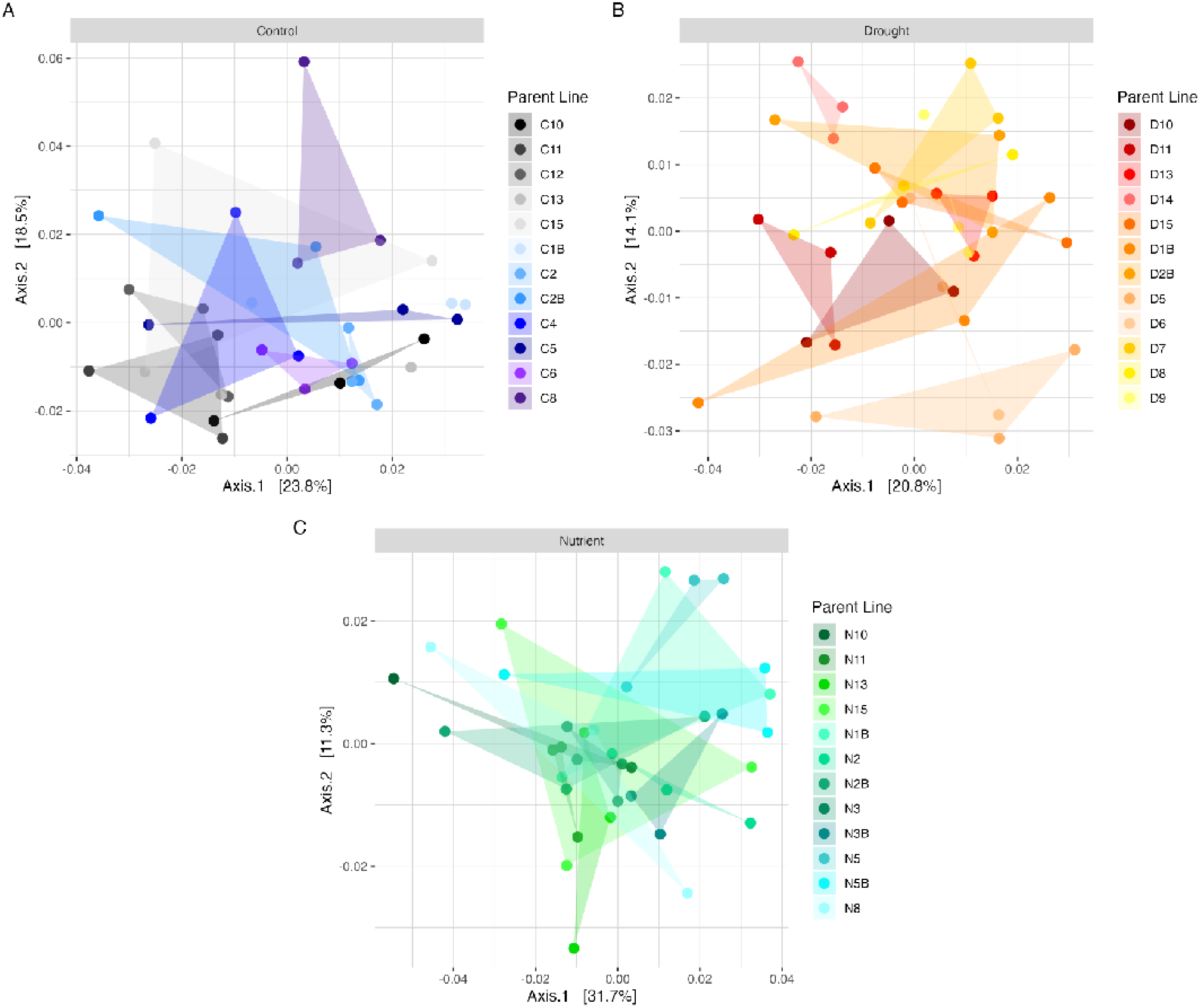
PCoA plots of Weighted Unifrac beta diversity in Generation 2, subset by parent plant treatment. When plant lines are grouped by G1 parent treatment, parent plant line is not significant in the control and nutrient lines, but is significant in the drought lines (PERMANOVA, Control: r^2^= 0.35319, F=1.2189, p=0. 0526, Nutrient: r^2^= 0.34740, F=1.1142, p= 0.2687, Drought: r^2^= 0.40311 F=1.4637 p=0.0021**). Bars above the ordinations indicated the treatment applied to the parent plants in G1: **A,** Control, **B,** Drought, **C,** Nutrient. The three offspring from each parent are connected by triangles to aid in visualization. Sample G2_9, the line C13 Nutrient offspring, was removed from figure A as an outlier. However, statistics were performed with this sample included.

**Fig. S5.**
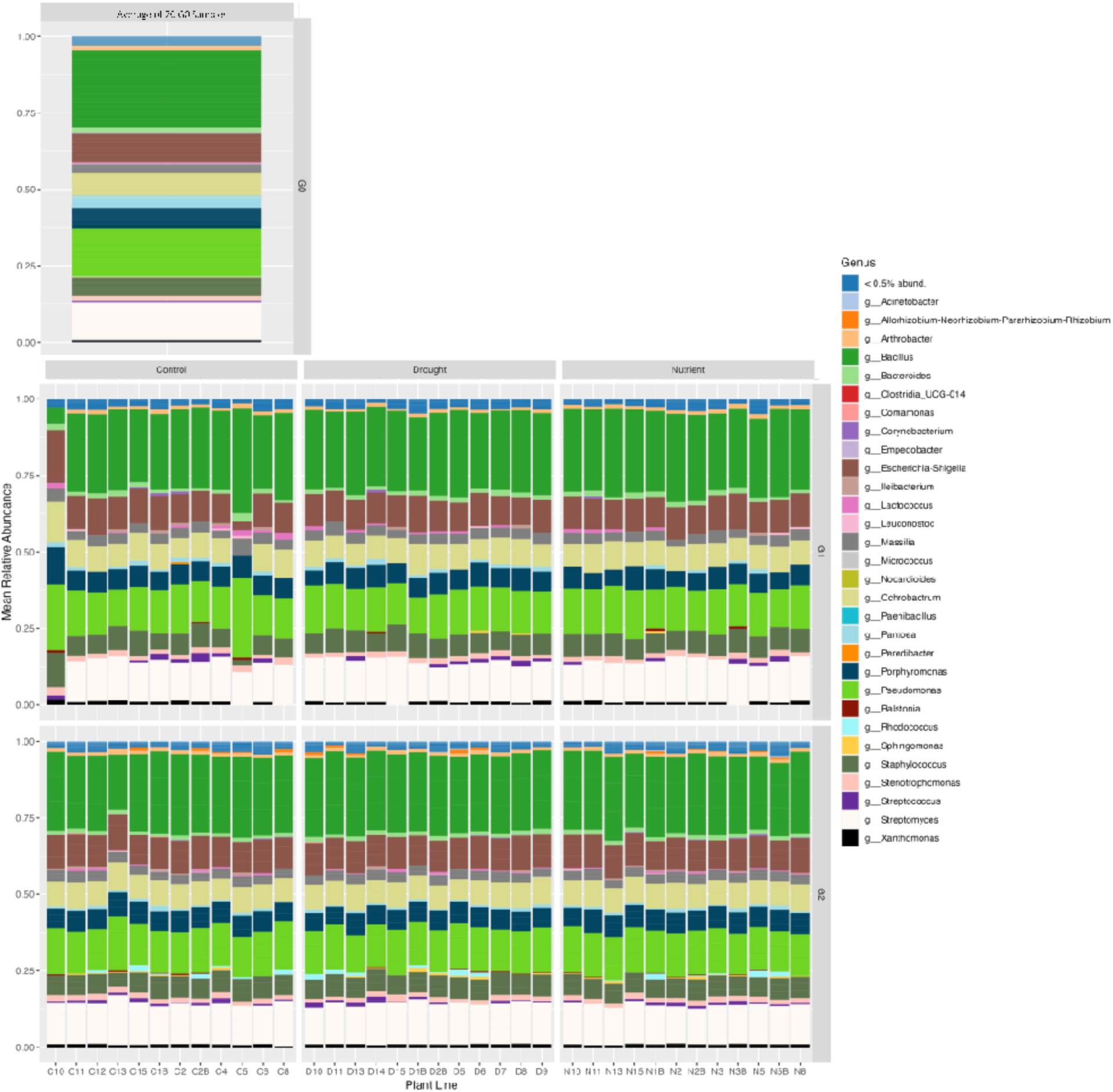
Mean relative abundance of full dataset ASVs identified at the genus level across all three generations. The bar in the top row is the average of the 20 seed samples from G0. Bars in the middle row represent G1 parent samples. The bars in the bottom row represent the average of the 3 G2 offspring samples in each parent line. The “< 0.5% abund.” category comprises 263 genera less than 0.5% abundant in the dataset. “g” indicates genus-level taxonomy.

**Fig. S6.**
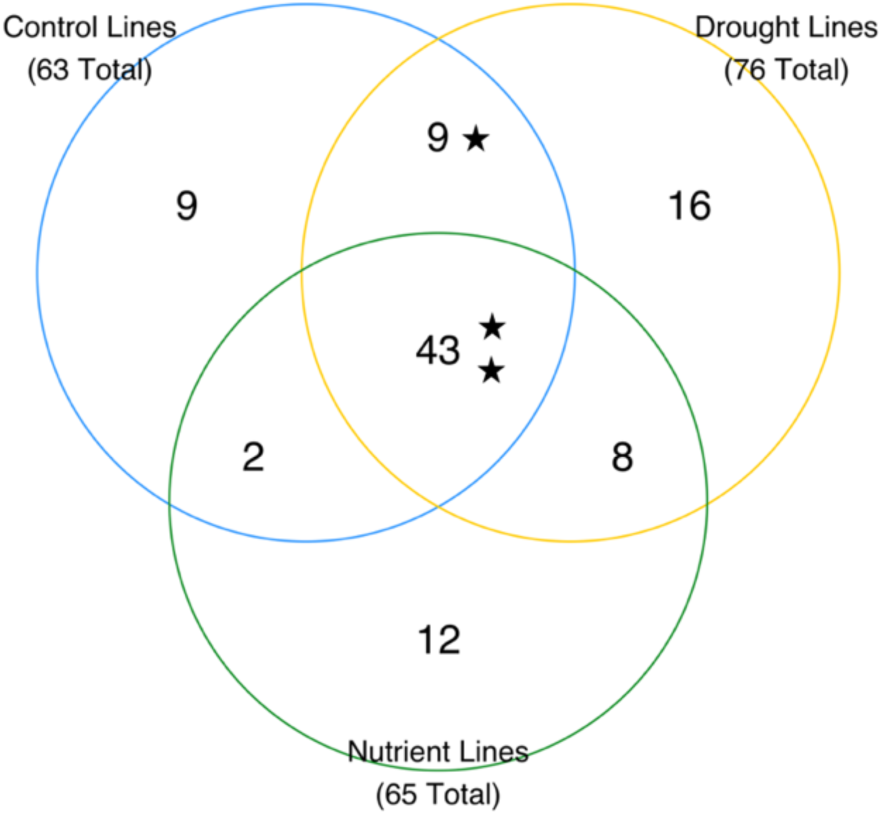
Number of ASVs found overlapping between G1 and G2 within parent lines and shared between parent treatment groups. Stars indicate the presence of core seed microbiome taxa identified by Simonin et al. 2022. There are 99 total ASVs represented, of which 85 can also be found in G0.

**Fig. S7.**
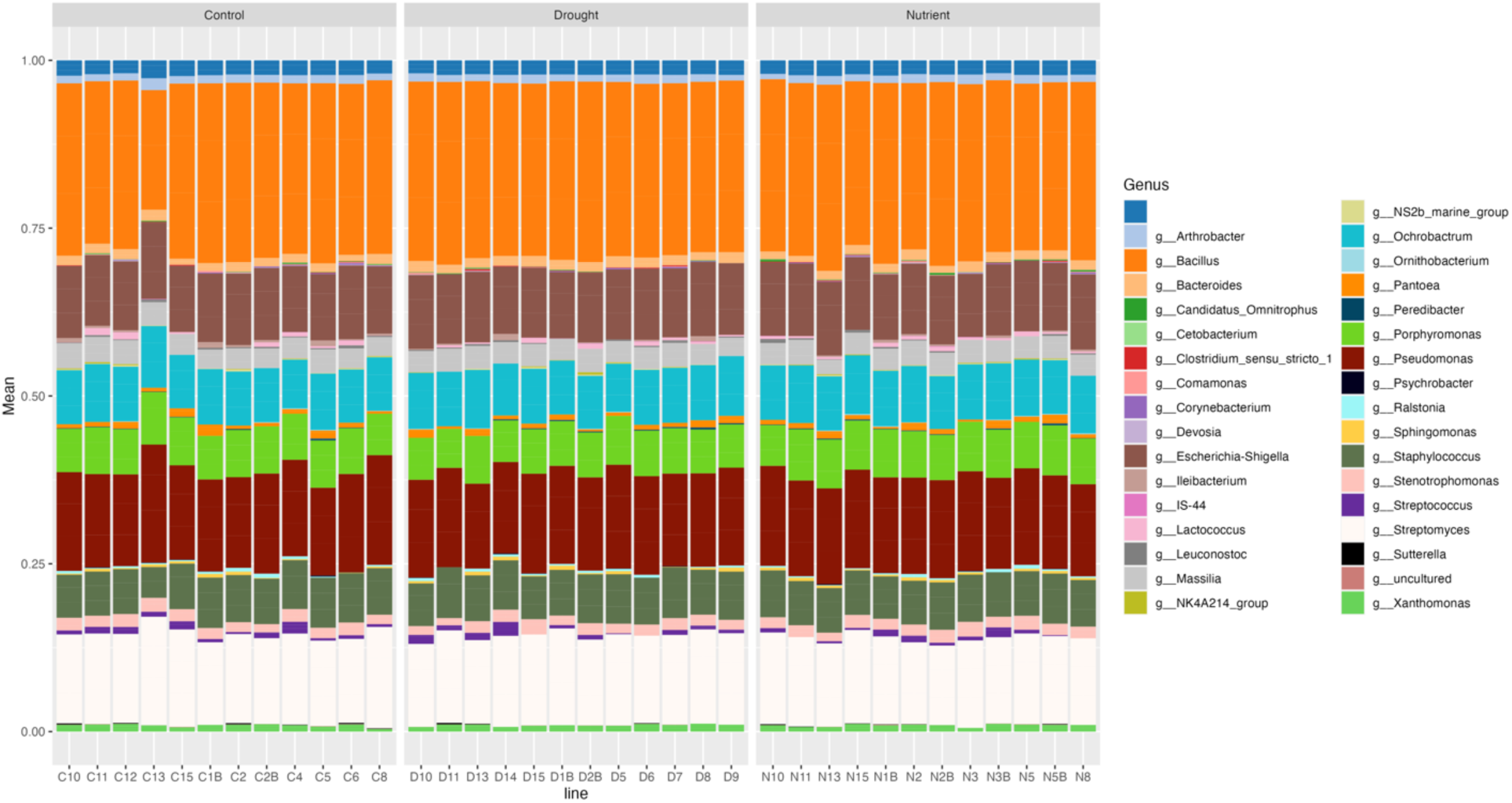
Mean relative abundance of 70 prevalent overlapping ASVs across parent lines in G2 identified at the genus level. All other ASVs have been removed from the dataset. The unlabeled legend color represents ASVs that are unresolved at the genus level.

**Fig. S8.**
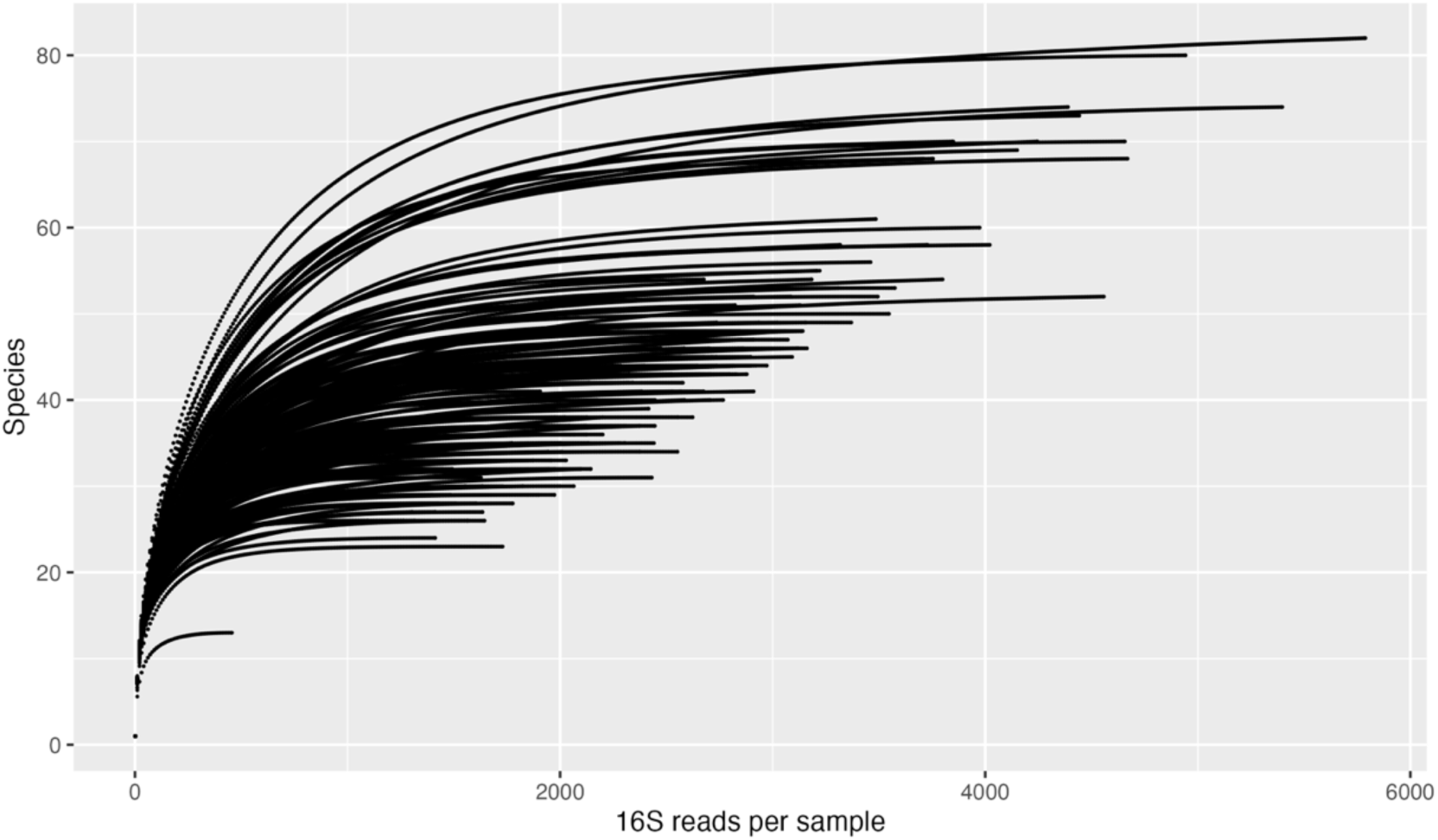
Rarefaction curves of quality-filtered microbiome profiles. (host reads removed, see methods). Each line represents one seed microbiome sample (pool of 5 seeds from the same parent plant). The DNA read range is 456-5788. All samples reached a plateau indicating a sufficient coverage to characterize seed microbiome diversity.

**Table S1.**
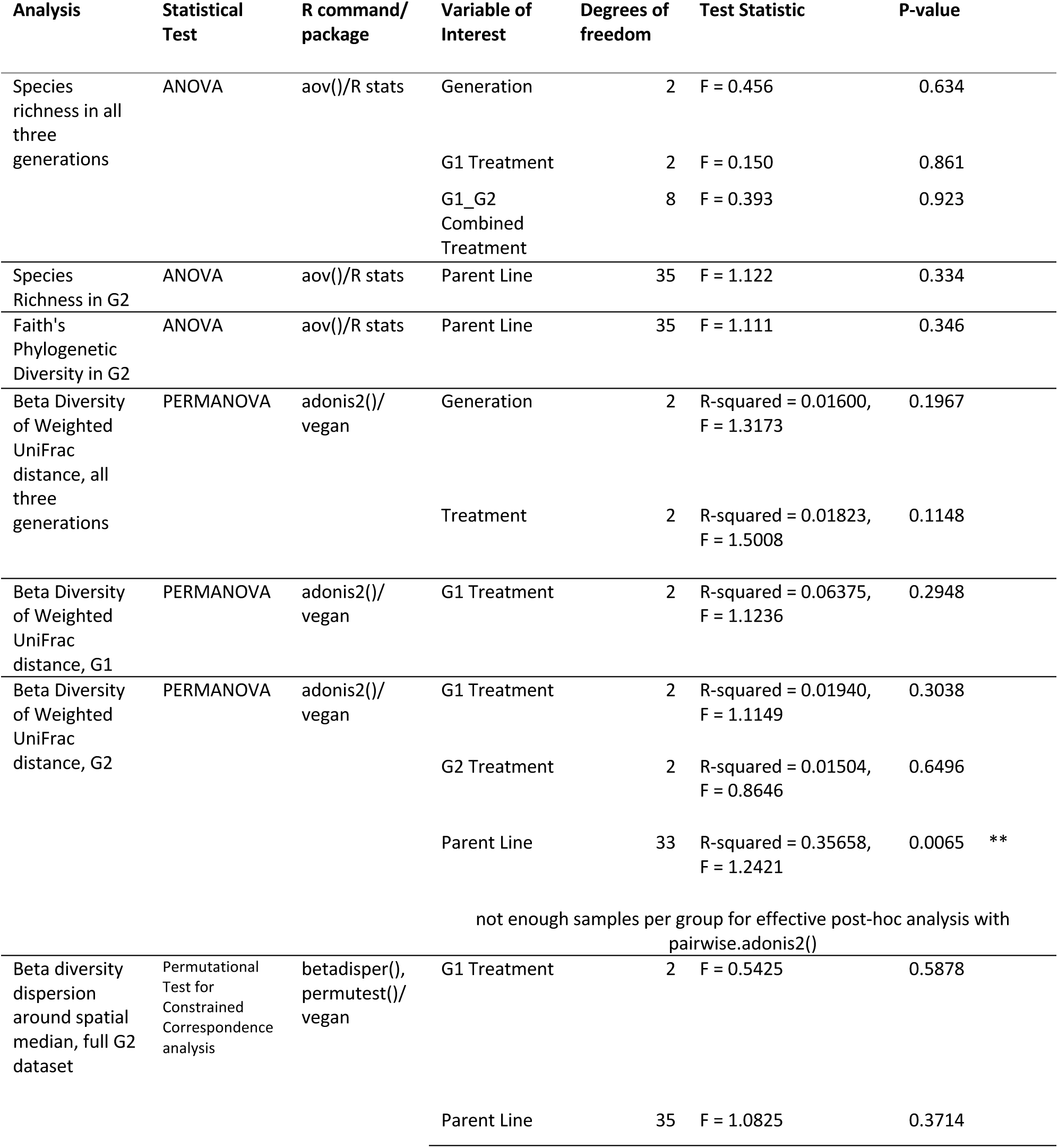
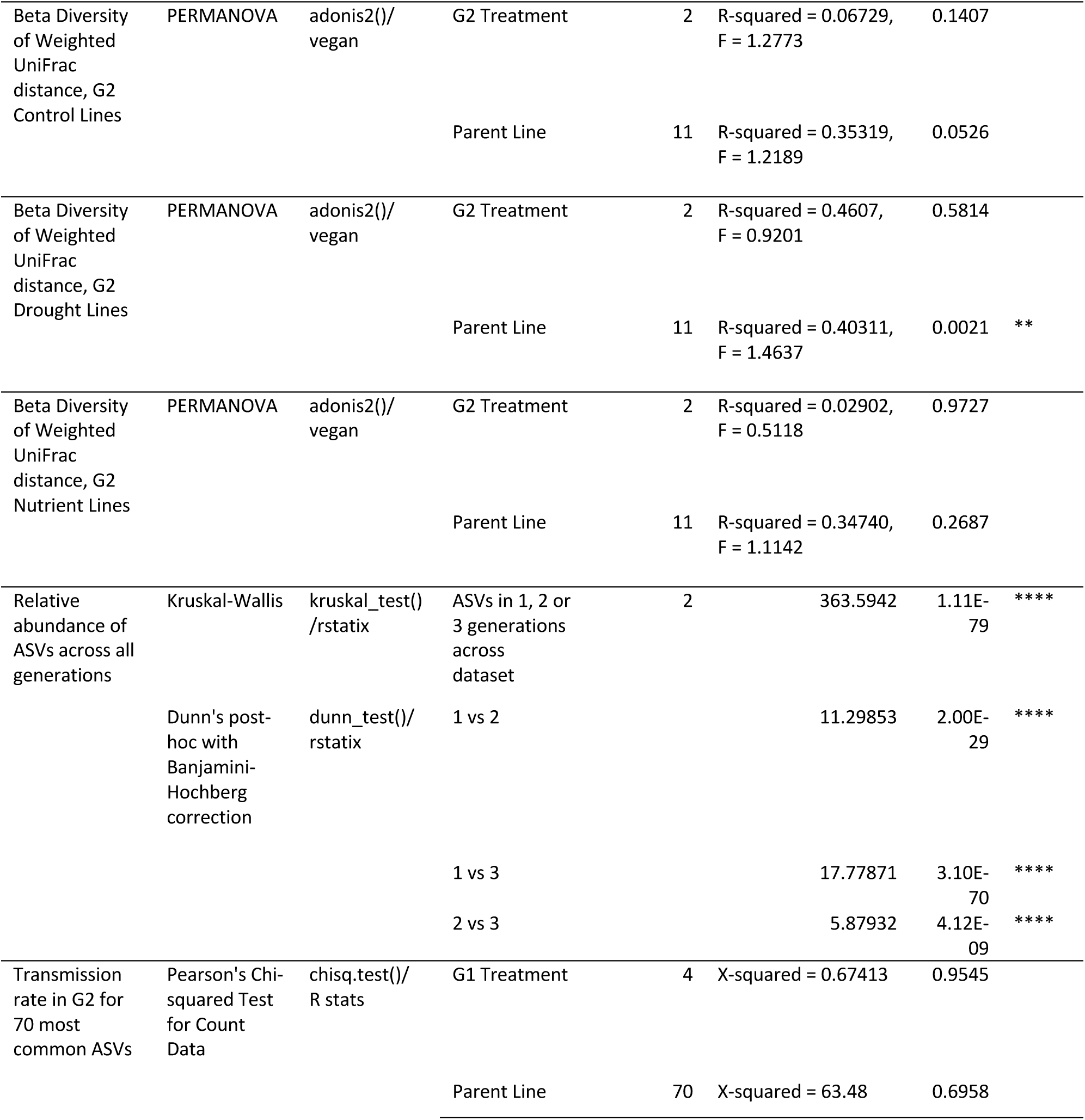
Statistical tests and their results to address hypotheses of differences in the alpha and beta diversity of the endophytic seed microbiome according to environmental treatment, parental line, plant generation, and their interactions.

**Table S2.** Excel file: Metadata table with MIMARKS compliant contextual information for each sample (e.g., plant treatment data, extraction batches).

**Table S3.** Excel file: List of 70 ASVs from Figure 2 with taxonomic identification, core taxa membership and % identity, and 16S V4 sequence.

